# Trends in *Arabidopsis* Research post genome sequencing- A Scientometric study

**DOI:** 10.1101/2022.02.22.481563

**Authors:** Sandeep Kumar, Amar Kant Kushwaha, R Thribhuvan, M Balakrishnan, P Krishnan

**Author notes:** **Corresponding authors** Sandeep Kumar & Krishnan P, Mail;, ICAR-National Academy of Agricultural Research and Management, Hyderabad, Telangana, India-500030.

## Abstract

*Arabidopsis thaliana*, a model plant, is intensively researched because of the intrinsic advantages associated with its life cycle, genetics, and other characteristics. In the present study, we analysed the publication data of research work done on *A. thaliana* during 2001-2020, a period when the whole genome sequence of the plant was available to the researchers. The current meta-analysis showed that 31965 research papers were published globally on *Arabidopsis* during the study period. The USA topped the countries with the maximum share of publications (28.97%), followed by China, Japan, and West European countries. The analysis showed a temporal shift in the number of publications from different countries and their research focus. After 2013, the research output from China was higher than that from the USA, and there was a shift in focus from developmental biology to stress-related topics. Plant Physiology journal carried the most research work (2490 publications) on Arabidopsis, which was followed by Plant Cell (2376), the Plant Journal (2260) and the Journal of Experimental Botany (1121). However, there was a progressive decline in the publication in these journals, and a shift was evident in favour of open access journals like Frontiers in Plant Science. The publication and citation numbers also show ongoing Arabidopsis research’s relevance to plant sciences, particularly curiosity-driven and discovery-based science. This article delves into the patterns in the prominence of research areas, ideas and foci for the future Arabidopsis research roadmap.

## 2. Introduction

*Arabidopsis thaliana* is a plant native to the Eurasian and African region, where it is generally considered a weed that often grows along the roadsides. Researchers across the globe have used this plant to study basic plant biology. It was first described by a German physician named Johannes Thal, on whose name Carl Linnaeus named it *Arabis thaliana*. The name *Arabidopsis thaliana* was given by Gustav Heynhold, who created a new genus by the name of *Arabidopsis* (Meyerowitz, 2001). It is self-pollinated, annual and grows up to 20-25cm. The key qualities, including small plant size, a relatively shorter life cycle period, seed setting in large quantities and smaller genome size with fewer chromosome number, qualifies it as the best candidate as a model plant and amenable for genetic and physiological studies (Rédei, 1970; Somerville and Koornneef, 2002; Koornneef and Meinke, 2010). Primary emphasis was given to the screening and identification of auxotrophic mutants (Rédei, 1975), as a result of which the first Arabidopsis gene sequences came out in 1986 (Chang and Meyerowitz, 1986).

After that, T-DNA-mediated transformation and vacuum infiltration made it more amenable to studies (Koncz et al., 1989). Eventually, Arabidopsis became a plant amenable to generating gene mutations and implementing DNA transformation protocols. The culmination of these projects was the National Science Foundation (NSF) project on Arabidopsis gene identification and whole-genome sequencing (Arabidopsis Genome Initiative, 2000). This project was one of many initiatives of genome sequencing. Various technological advancements happened after the completion of Arabidopsis genome sequencing, which shaped the biological research in many organisms. Genome sequencing brought a tremendous pace and shift in Arabidopsi*s* research (Bevan and Walsh, 2005). Multinational Arabidopsis Steering Committee (MASC) has been actively involved in Arabidopsis research and constantly prepares the roadmaps for Arabidopsis research (http://arabidopsisresearch.org/index.php/en/about). The research interest in *Arabidopsis* as a model plant in the emerging research scenarios has been dwindling following the emergence of high throughput sequencing and other genomics tools (Fig. 1). In addition, there has been an increased application of these newer tools to non-model plants. The current research was designed to analyse the trends in research on the *Arabidopsis thaliana*, in the post-genome sequencing period, with the specific objectives of

**Fig. 1:**
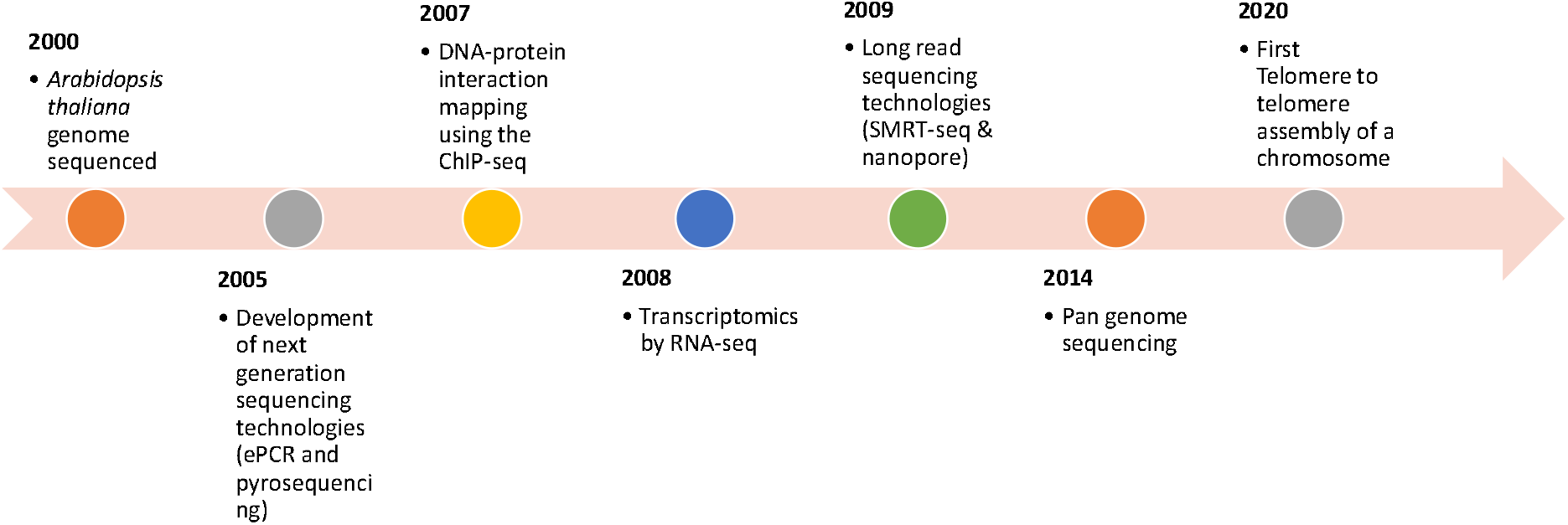
Timeline of critical technological advancements in Genome studies after *A. thaliana* genome Sequencing. (Adapted from *Nature Milestones: Genomic Sequencing* https://www.nature.com/collections/genomic-sequencing-milestones (2021))

i. Mapping the variation in the theme areas of Arabidopsis research and
ii. Profile the trends in research networks at various levels, *viz*., authors, organisations and countries.

## 3. Materials and methods

### 3.1 Data collection and analysis

This study uses the bibliometric methodology to quantify scientific collaborations in Arabidopsis research worldwide. The Web of Science (WoS) database was used for the literature search on Arabidopsis. Web of Science is a website published by Clarivate Analytics that offers subscription-based access to numerous databases that provide detailed citation data for many academic subjects (Li et al., 2018). The search string and filters used to select the publications were as follows (T.I. = (Arabidopsis)) AND DOCUMENT TYPES: (Article) Timespan: 2001-2020. Indexes: SCI-EXPANDED, CPCI-S, ESCI. The year 2001 was chosen as the starting point for the survey since Arabidopsis genome sequencing had been finished by 2000 (Arabidopsis Genome Initiative, 2000). Conference proceedings and books/book chapters were excluded from the study. The search phrase was chosen based on our investigation of existing literature and discussions among authors and subject matter experts. The *thaliana* was not added to the string as our preliminary scanning showed that many publications did not use the full binomial name in the titles and used only ‘*Arabidopsis*’.

The data retrieved from the WoS database was exported in text format, with each record containing the bibliometric data: title name, journal names, author names with Web of Science affiliation, number of citations, document kinds, year of publication for each article, authors, Web of Science keywords, and abstracts. The WoS database was also utilised for publication trends over the years, for which five year moving average of publications was used. The Scientometric/bibliometric parameters like the total number of publications and citations, average citations per article (ACPP), h-index, research areas, collaborating countries and funding agencies were extracted from the WoS platform for 20 years (2000-2020).

### 3.2 Network analysis

The co-authorship and keyword occurrence maps were generated using the VOS viewer version 1.6.16 using metadata generated from the web of science (van Eck and Waltman, 2011) (http://www.vosviewer.com/). For the network analysis, the top 100 results were selected and used in case of both institutional collaboration and year wise keyword occurrence. Bibliometrix, an R-based tool, was used for data filtering, research theme evolution and identifying journals based on Bradford’s law, wherein the metadata was fed in BibTeX files (http://www.bibliometrix.org) (Aria and Cuccurullo, 2017).

## 4. Results & Discussion

### 4.1 Global output of research on *Arabidopsis*

The publication of the *Arabidopsis* genome led to a significant rise in *Arabidopsis* research as it expanded the area of research from only isozyme and protein study to functional genomics (Mysore et al., 2001; Nath Radhamony et al., 2005; Bevan and Walsh, 2005). The study showed that 31965 research articles were published on the subject during the 20 years from 2001 to 2020 (Fig. 2). One intriguing element of the total number of articles was a five-year consecutive decline in the total number of publications following 2013, except for 2019/2020. This decreasing trend in recent years can be attributed to increased emphasis on the sequencing of commercially significant species such as rice, maise, pigeon pea, rapeseed, tomato and many others (International Rice Genome Sequencing Project, 2005; Schnable et al., 2009; Varshney et al., 2011; Wang et al., 2011). This paradigm shift was made possible due to the remarkable progress of sequencing techniques appropriate for diverse purposes, from high cost and poor throughput to low cost and high throughput (Türktaş et al., 2015).

**Figure 2:**
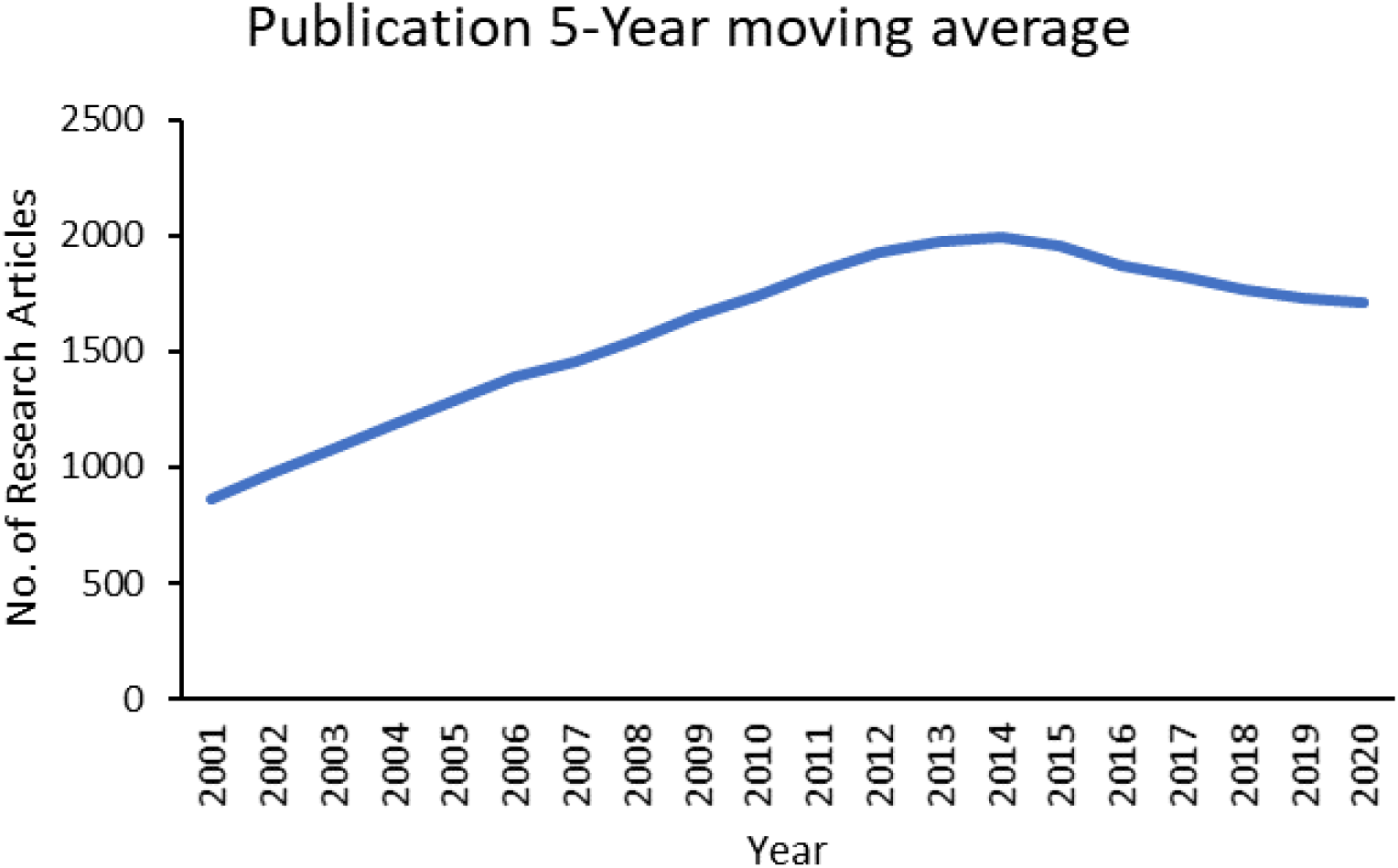
Research output in *Arabidopsis* over the years

In the keyword co-occurrence, we found a clear shift from developmental pathways related to the peripheral pathways like stress responses and the emergence of some new keywords like transcriptome and proteomics in the last decade (Fig. 3). It also demonstrates the widespread usage of various next-generation genomic tools and techniques in the previous decade (2011-2020). The *Arabidopsis* gene functions have also been elucidated in the major signalling pathways (Alonso and Stepanova, 2004; Atkinson et al., 2013). Bolker (2012) emphasised the need to go beyond model organisms to get a complete picture of various biochemical and signalling pathways in taxonomic groups. Moreover, the elucidation of biological function through *Arabidopsis* has trickled into the agriculturally important crop sp. for their validation and utilisation (Priyanka et al., 2010; Manmathan et al., 2013; Mishra et al., 2014).

**Figure 3:**
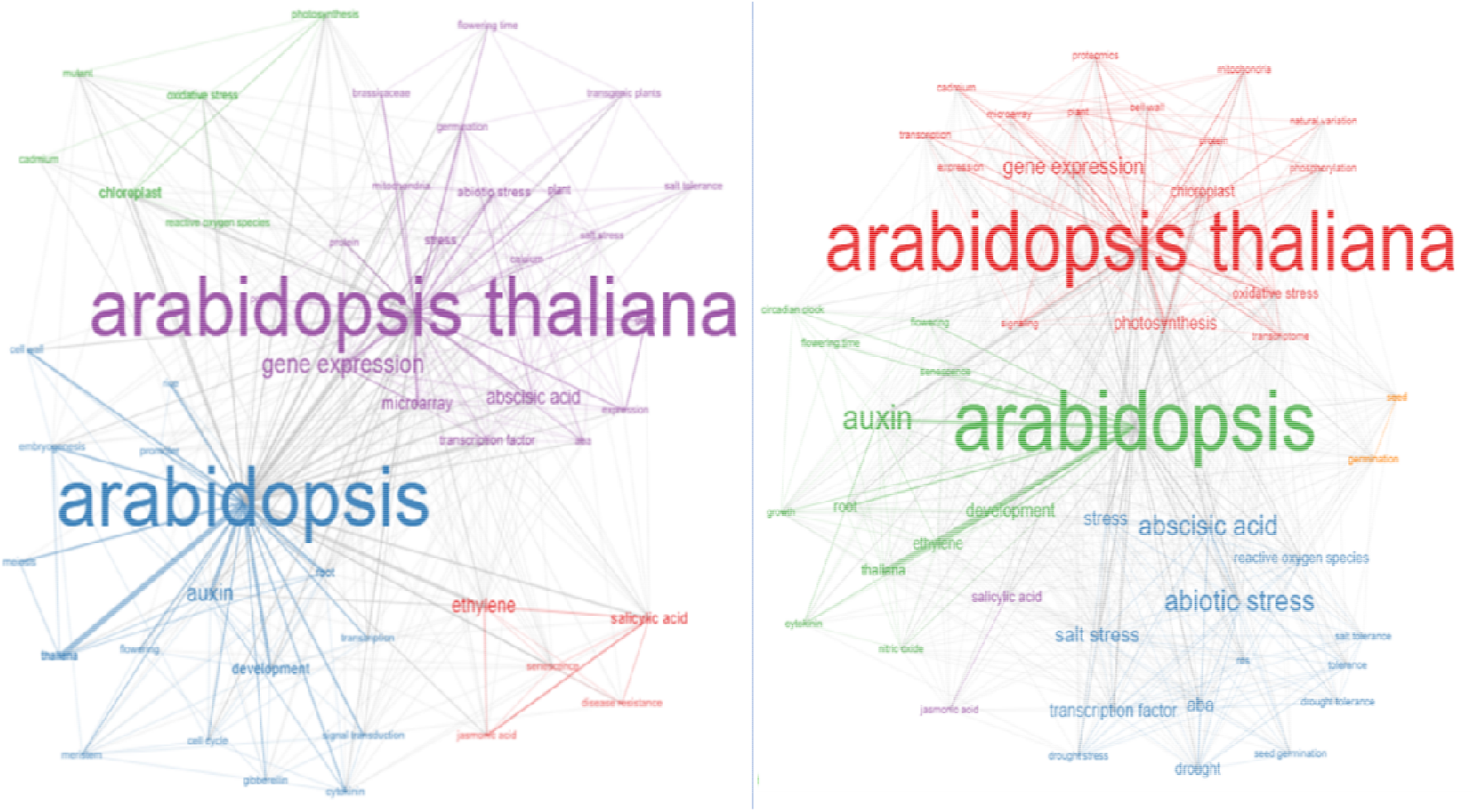
Thematic evolution based on the major keywords co-occurrence in two decades (2001-10) & (2011-20)

### 4.2 Highly cited studies on Arabidopsis

After whole-genome sequencing of Arabidopsis, the research papers on insertional mutagenesis in *Arabidopsis thaliana* received more citations, thus indicating the research trends (Table 1). Before sequencing, it was tough to create a loss of function mutation. Till the sequencing of the first plant, only unicellular organisms were subject to the creation of loss of mutation for understanding the function of the gene(s) (Giaever et al., 2002) and gene silencing and RNAi were utilised for understanding the developmental role of genes in *C. elegans* with their limitations of unstable heritability, residual expression of genes and low throughput (Ashrafi et al., 2003). After the publication of the paper titled “Genome-wide Insertional mutagenesis of *Arabidopsis thaliana”* by Alonso et al (2003) depicting the technique overcoming the limitation of earlier methods, work started in Arabidopsis mutant generation and their use in research. The article influenced a large stream of research, as indicated by the maximum citations (3790) that any Arabidopsis paper received in the last two decades and laid the foundations of functional biology in Arabidopsis.

**Table 1:** Highly cited research publications on *Arabidopsis* since its full genome was sequenced

| No | Title of the paper | Author and year | Affiliation of corresponding author | Journal | Citations (WoS) |
| --- | --- | --- | --- | --- | --- |
| 1. | Genome wide insertional mutagenesis of *Arabidopsis thaliana* | Alonso J.M. et al., 2003 | Salk Inst Biol Studies, Genome Anal Lab, 10010 N Torrey Pines Rd, La Jolla, CA 92037 USA. | Science | 3790 |
| 2. | *Arabidopsis* mesophyll protoplasts: a versatile cell system for transient gene expression analysis | Yoo S.D., Cho Y.H. and Sheen J. (2007) | Massachusetts Gen Hosp, Dept Mol Biol, Boston, MA 02114 USA. | Nature Protocols | 2344 |
| 3. | Genome-wide identification and testing of superior reference genes for transcript normalisation in *Arabidopsis* | Czechowski, T. et al., 2005 | Max Planck Inst Mol Plant Physiol, D-14476 Potsdam, Germany. | Plant Physiology | 2018 |
| 4. | GENEVESTIGATOR. *Arabidopsis* microarray database and analysis toolbox | Zimmermann P. et al., 2004 | ETH, Inst Plant Sci, CH-8092 Zurich, Switzerland. | Plant Physiology | 1888 |
| 5. | A gene expression map of *Arabidopsis thaliana* development | Schmid, M. et al., 2005 | Max Planck Inst Dev Biol, Spemannstr 37-39, D-72076 Tübingen, Germany | Nature Genetics | 1787 |
| 6. | MAP kinase signalling cascade in *Arabidopsis* innate immunity | Asai, T. et al., 2002 | Harvard Univ, Sch Med, Dept Genet, Boston, MA 02114 USA. | Nature | 1684 |

Similarly, with the advancement in sequencing techniques, epigenomes were targeted to decipher the developmental pathway. To detect such epigenetic modification, Cokus et al (2008) came up with a new approach, BS-seq (Bisulphite sequencing), combining sequence information and bisulphate treatment for mapping the methylation sites in the whole genome. Similarly, in the same area, Lister et al (2008) mapped the methylation sites by sequencing smRNA, and methylcytosine to decipher the role in the developmental pathway. The significance of these two articles is indicated by the citations received by them, viz., 1516 and 1363, respectively. Some other papers featured on the top 10 list by the number of citations were related to transcript normalisation, transient gene expression and microarray database. In short, the protocol and database development paper featured prominently on the most cited list. It is in line with the evolution of *Arabidopsis* as a model plant, evidenced by the fact that novel tools and techniques to decipher plant biological functions has been developed in Arabidopsis and later extended to other crop plants (Koornneef and Meinke, 2010).

### 4.3 Top countries working on the Arabidopsis research

Based on the number of publications, America contributed 28.97% of the total publications, followed by China, Germany, and Japan with 21.23%, 14.11% and 12.06% of the publications, respectively. Table 2 shows the top 10 countries and their per cent contribution, along with other publication metrics *viz*., h-index, T.C. (Total Citations) and ACPP (Average Citation Per Publication). The h-index, which indirectly represents the research influence, was higher for publications from the USA (133), which was followed by Germany (223), England (204) and Japan (200). Though China has the second-highest per cent share in total research papers, its h-index is relatively lower, i.e., 166 and the lowest ACPP score of 29.96 compared to other top 10 countries. The country collaboration map shown in Fig. 5 provides a clear picture of the collaborations. The United States of America and the countries in Western Europe like the UK, France and Germany showed a very high collaboration and scientific output. However, some African and Baltic countries were missing from the Arabidopsis research map entirely. The countries leading the research in terms of the number of publications were part of the genome sequencing consortium or those collaborating with them (https://www.nsf.gov/bio/pubs/reports/arab_prog78.htm). Though China was not the part as they were not much into *Arabidopsis* research before 1995 (Chen et al., 2006), its research output has been high in recent years (Fig 4). A plausible explanation of this is the acquisition of natural and mutant stocks of Arabidopsis seeds post-genome sequencing (Provart et al., 2016).

**Table 2:**
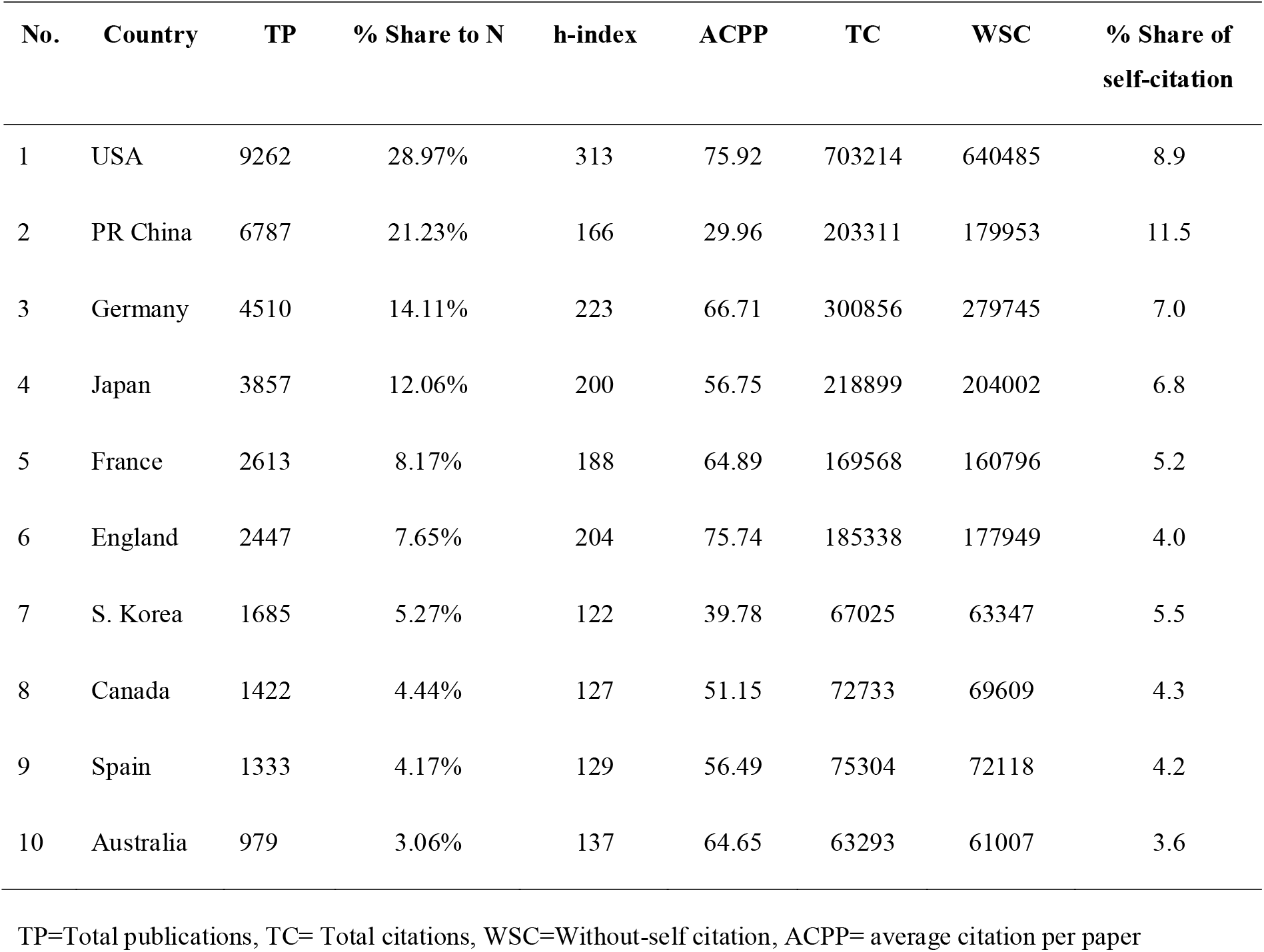
Top 10 countries which published the most research papers on *Arabidopsis* in the world during 2001-2020

| No. | Country | TP | % Share to N | h-index | ACPP | TC | WSC | % Share of self-citation |
| --- | --- | --- | --- | --- | --- | --- | --- | --- |
| 1 | USA | 9262 | 28.97% | 313 | 75.92 | 703214 | 640485 | 8.9 |
| 2 | PR China | 6787 | 21.23% | 166 | 29.96 | 203311 | 179953 | 11.5 |
| 3 | Germany | 4510 | 14.11% | 223 | 66.71 | 300856 | 279745 | 7.0 |
| 4 | Japan | 3857 | 12.06% | 200 | 56.75 | 218899 | 204002 | 6.8 |
| 5 | France | 2613 | 8.17% | 188 | 64.89 | 169568 | 160796 | 5.2 |
| 6 | England | 2447 | 7.65% | 204 | 75.74 | 185338 | 177949 | 4.0 |
| 7 | S. Korea | 1685 | 5.27% | 122 | 39.78 | 67025 | 63347 | 5.5 |
| 8 | Canada | 1422 | 4.44% | 127 | 51.15 | 72733 | 69609 | 4.3 |
| 9 | Spain | 1333 | 4.17% | 129 | 56.49 | 75304 | 72118 | 4.2 |
| 10 | Australia | 979 | 3.06% | 137 | 64.65 | 63293 | 61007 | 3.6 |

TP=Total publications, TC= Total citations, WSC=Without-self citation, ACPP= average citation per paper

**Figure 4:**
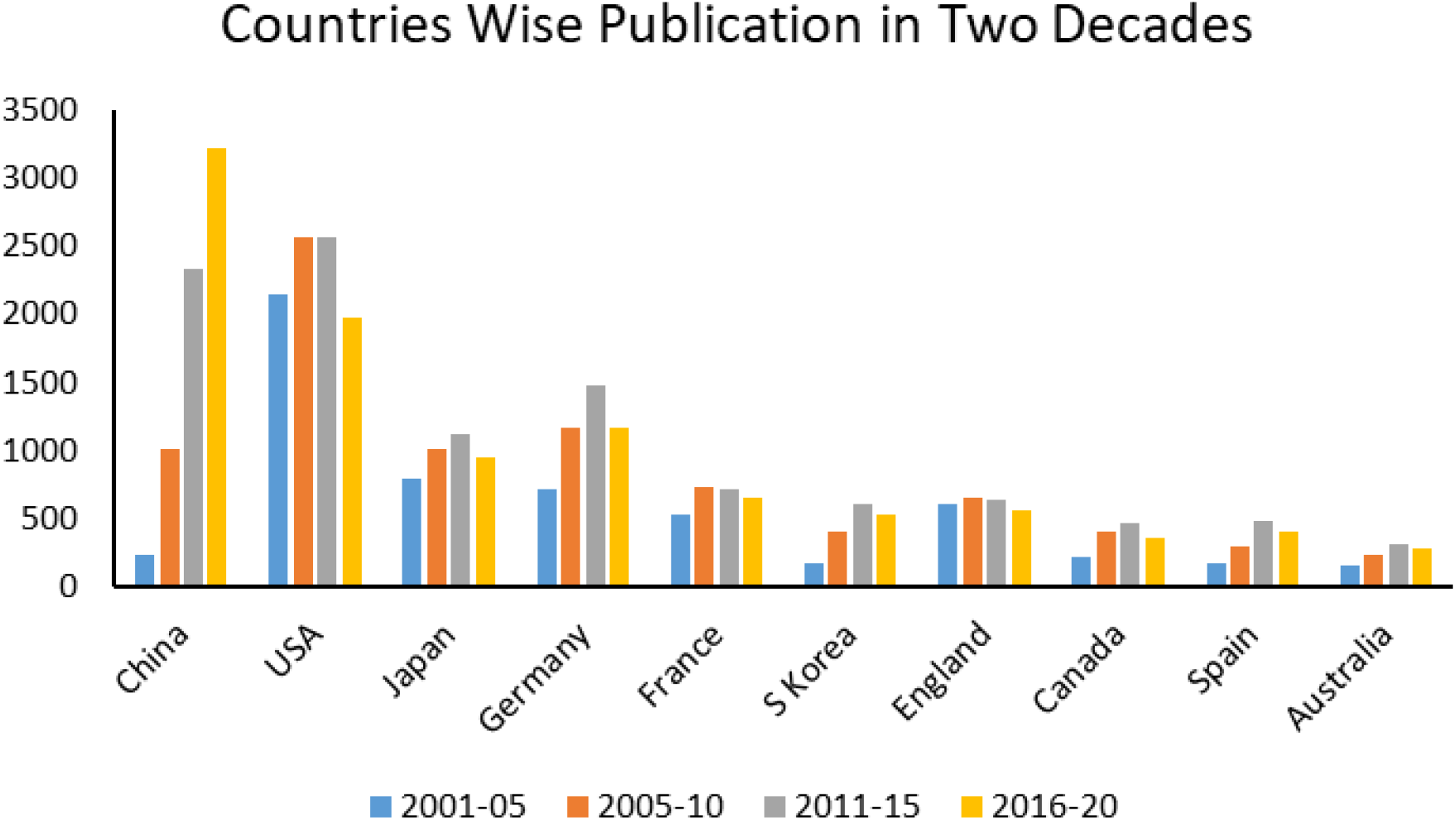
Country-wise publication over the different period

**Figure 5:**
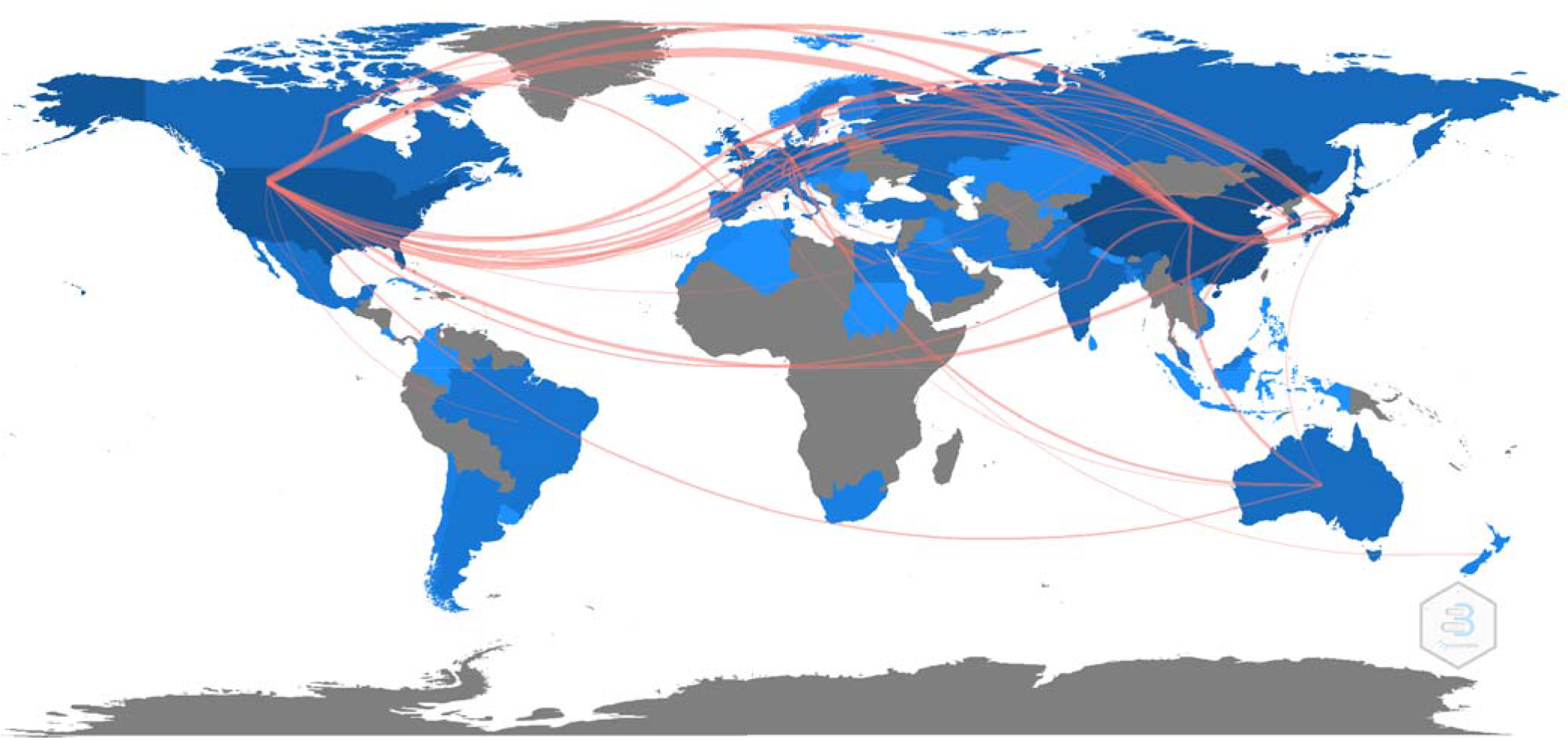
Country-wise Scientific production based on the number of publications and Interaction among countries publishing articles on *A. thaliana*. The blue colour’s intensity corresponds to the total number of publications from the corresponding country, and the red lines highlight the interactions among countries (co-authorship).

Based on subject areas, most publications featured the plant sciences and developmental biology across the countries (Table 3). Cell biology and developmental biology were laid less stress in China, possibly due to more research being done later. However, the research areas of publication journals are indicative, and there might exist many cross-cutting and multidisciplinary studies which do not follow this classification.

**Table 3:** Top 10 Country vs Subject field wise (No. of publications)

| Country/R<br>egion | Subject |  |  |  |  |  |  | Total* |
| --- | --- | --- | --- | --- | --- | --- | --- | --- |
|  | Plant<br>Sciences | Bio-<br>chemistry<br>and<br>Molecular<br>biology | Cell<br>Biology | Muti-<br>disciplinary<br>Sciences | Genetic<br>heredity | Bio-<br>technology<br>applied<br>micro-<br>biology | Developm<br>ental<br>biology |  |
| USA | 5549 | 3014 | 1760 | 1053 | 795 | 419 | 243 | 9262 |
| China | 4351 | 1976 | 753 | 616 | 410 | 361 | 55 | 6787 |
| Germany | 2885 | 1479 | 857 | 407 | 313 | 131 | 106 | 4510 |
| Japan | 2559 | 1206 | 1031 | 285 | 194 | 366 | 75 | 3857 |
| France | 1583 | 885 | 507 | 223 | 193 | 86 | 59 | 2613 |
| England | 1495 | 770 | 477 | 257 | 158 | 88 | 66 | 2447 |
| S. Korea | 1080 | 595 | 320 | 96 | 65 | 110 | 17 | 1685 |
| Canada | 1011 | 367 | 191 | 89 | 114 | 94 | 30 | 1422 |
| Spain | 880 | 392 | 194 | 133 | 82 | 57 | 40 | 1333 |
| Australia | 689 | 251 | 149 | 90 | 45 | 28 | 18 | 979 |

\*The total number of articles in sub-subjects does not add up as only the five major subjects are mentioned here, and there exist many journals which fall under more than one category

### 4.4 Leading institutions in the Arabidopsis research

Collaborations between different institutes revealed the close links between American and European institutes (Fig. 5). The top three institutions with the most publications were the Chinese Academy of Science, the University of California, and the University of Tokyo. However, the Asian institutes had relatively weak partnerships with other countries. The North American and European institutes were active in the early research periods, as visible from the corresponding colour. The Chinese Academy of Science and other Chinese Universities have published more papers than those from Japan and Western countries.

The highest papers feature the Chinese Academy of Sciences (Fig. 6). However, the articles from western Europe had relatively high-quality parameters, e.g. Chinese Academy of science has 1856 total publications, but the average citation per paper is lower (41.69) than the U.K. Research Innovation (UKRI). UKRI has only 779 total papers but 90.03 average citations per paper.

**Figure 6:**
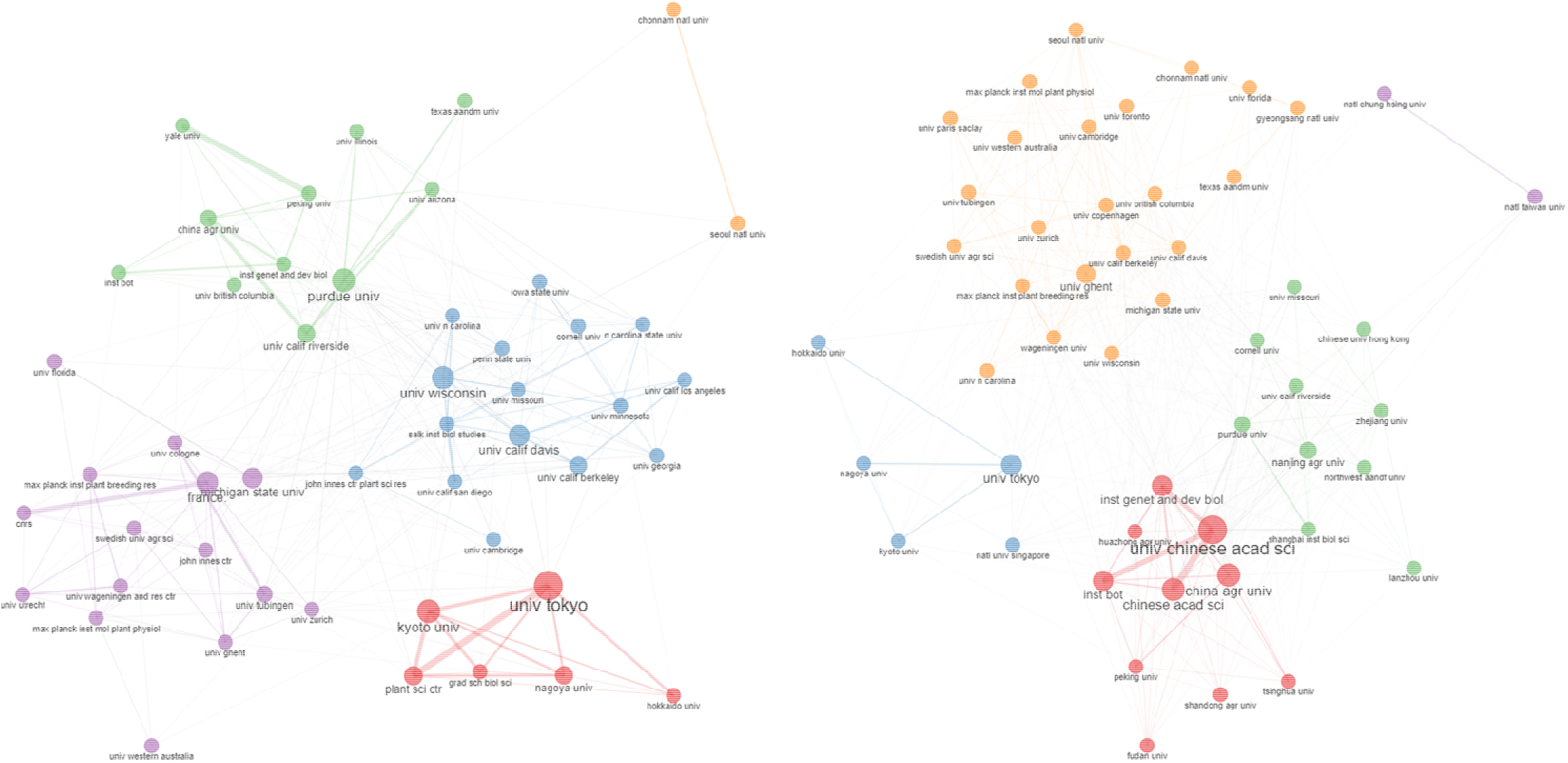
Collaboration among different institutions working on *Arabidopsis* in the first decade (2001-2010) vs second decade (2011-2020)

### 4.5 Top Journals for Arabidopsis Research

Plant Physiology (2376), Plant cell (2490) and the Plant Journal (2260) were the top three journals in terms of the number of relevant publications, h-index, and total citations (Table 4). Bradford law is an essential consideration that decides the importance of different journals based on diminishing citation of relevant articles over the year, following Pareto distribution (Nash-Stewart et al., 2012). Here it can be easily seen that Plant Physiology, Plant Cell, Plant Journal, Journal of Experimental Botany, Plant and Cell Physiology, Frontiers in Plant Science and PNAS were the core sources that carried most publications on Arabidopsis and thus represent the choice journals for the researchers (Fig. 6). The annual pattern of publication on Arabidopsis in these journals showed a shift in the preferred journals over the years (Fig. 7), i.e. a decline in the publication in the Plant cell, plant physiology and the plant journal; however, open publications models of frontiers in plant sciences have gained in popularity. As a result, none of the top three journals based on the total number of publications was present among the top three for the recent year, i.e. 2020. This trend is probably due to the citation advantage of an article in open access (O.A.) due to an increase in the number of readers citing open access articles (Cullen and Chawner, 2011). The radical initiative to O.A. rather than a subscription with ‘Plans S’ (science, speed, solution, shock) by 11 funding agencies in Europe indicates such implication (Else, 2018).

**Table 4:** Top journals that published most *Arabidopsis* research articles from 2001 to 2020

| No. | Title of the journal | TP | % share | TC | WCS | H-Index | ACPP | IF (2020) |
| --- | --- | --- | --- | --- | --- | --- | --- | --- |
| 1 | *Plant Physiology* | 2490 | 7.79 | 226960 | 220788 | 204 | 91.15 | 6.902 |
| 2 | *Plant Cell* | 2376 | 7.43 | 287284 | 277416 | 248 | 120.21 | 9.618 |
| 3 | *Plant Journal* | 2260 | 7.07 | 179334 | 174911 | 190 | 79.35 | 6.141 |
| 4 | *Journal of Experimental Botany* | 1121 | 3.51 | 44776 | 44089 | 99 | 39.94 | 5.908 |
| 5 | *Plant and Cell Physiology* | 999 | 3.12 | 46314 | 45320 | 105 | 46.36 | 4.062 |
| 6 | *Frontiers in Plant Science* | 950 | 2.97 | 13016 | 12696 | 44 | 13.7 | 4.407 |
| 7 | *PNAS* | 878 | 2.75 | 102939 | 101908 | 180 | 117.24 | 9.412 |
| 8 | *PLoS ONE* | 829 | 2.59 | 23548 | 23269 | 67 | 28.41 | 2.740 |
| 9. | *Plant Signalling Behaviour* | 804 | 2.52 | 8810 | 8728 | 36 | 10.96 | 1.671 |
| 10. | *Plant Molecular Biology* | 677 | .12 | 33566 | 33234 | 90 | 49.58 | 3.302 |

**Figure 7:**
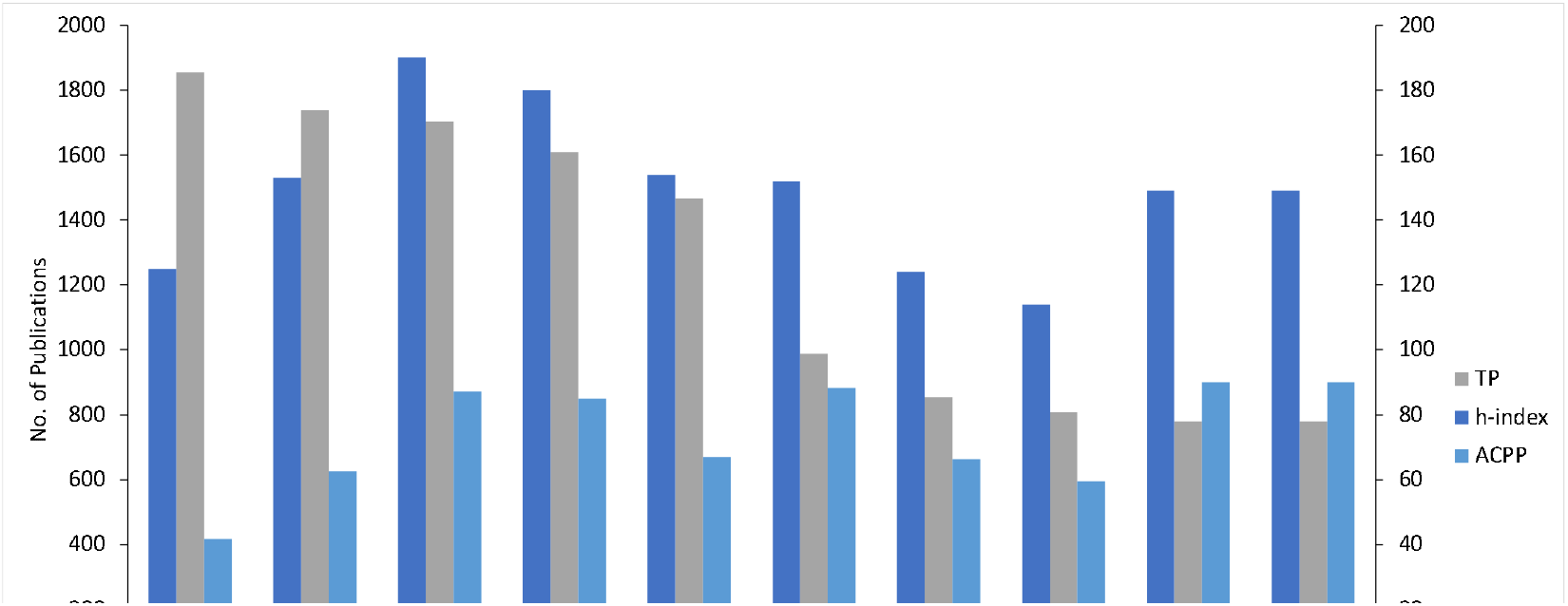
Quantitative and qualitative parameters in the research backed by different funding organisations (2001-2020)

**Figure 8:**
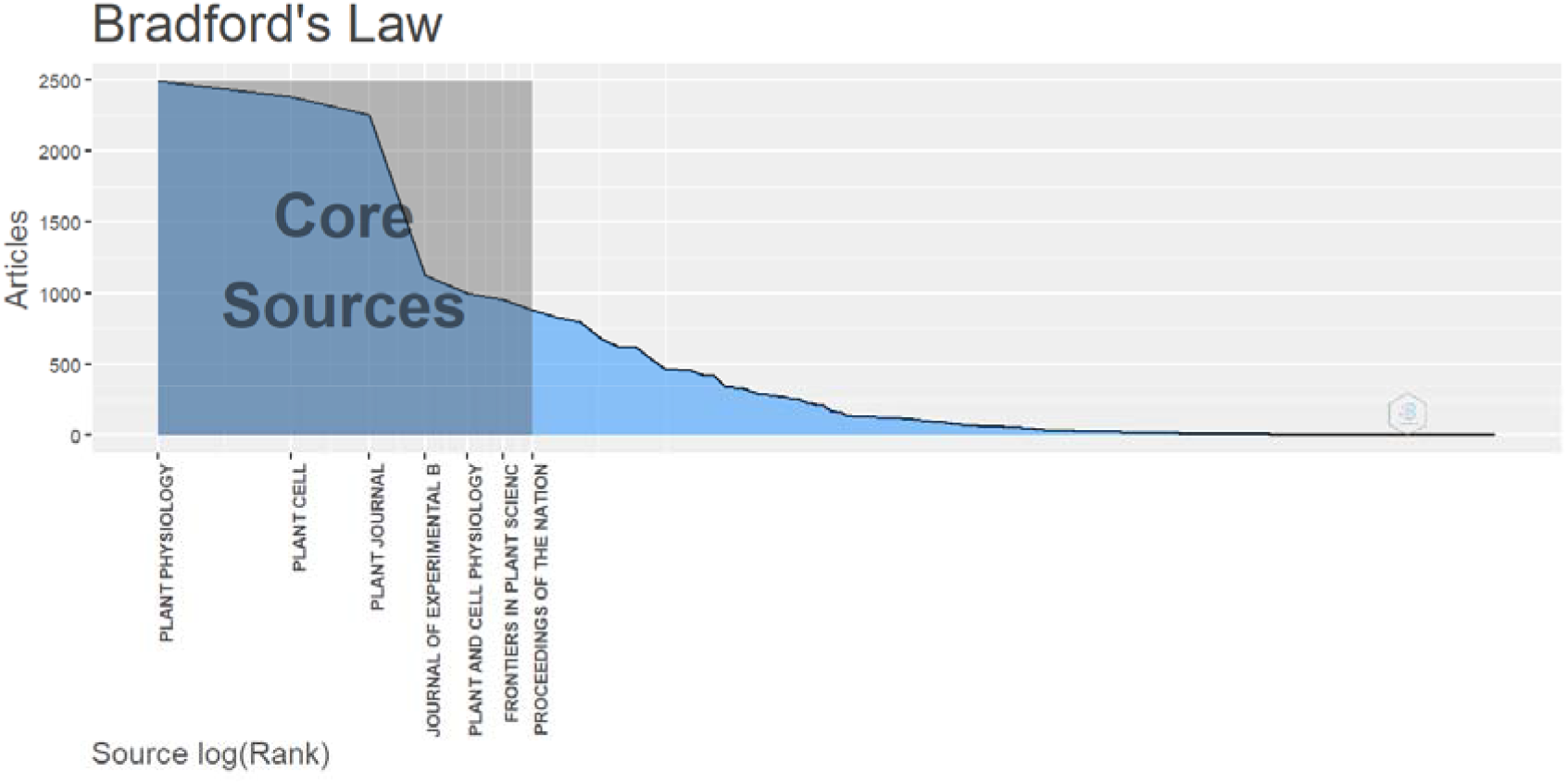
The Bradford law displaying key Arabidopsis journals

**Figure 9:**
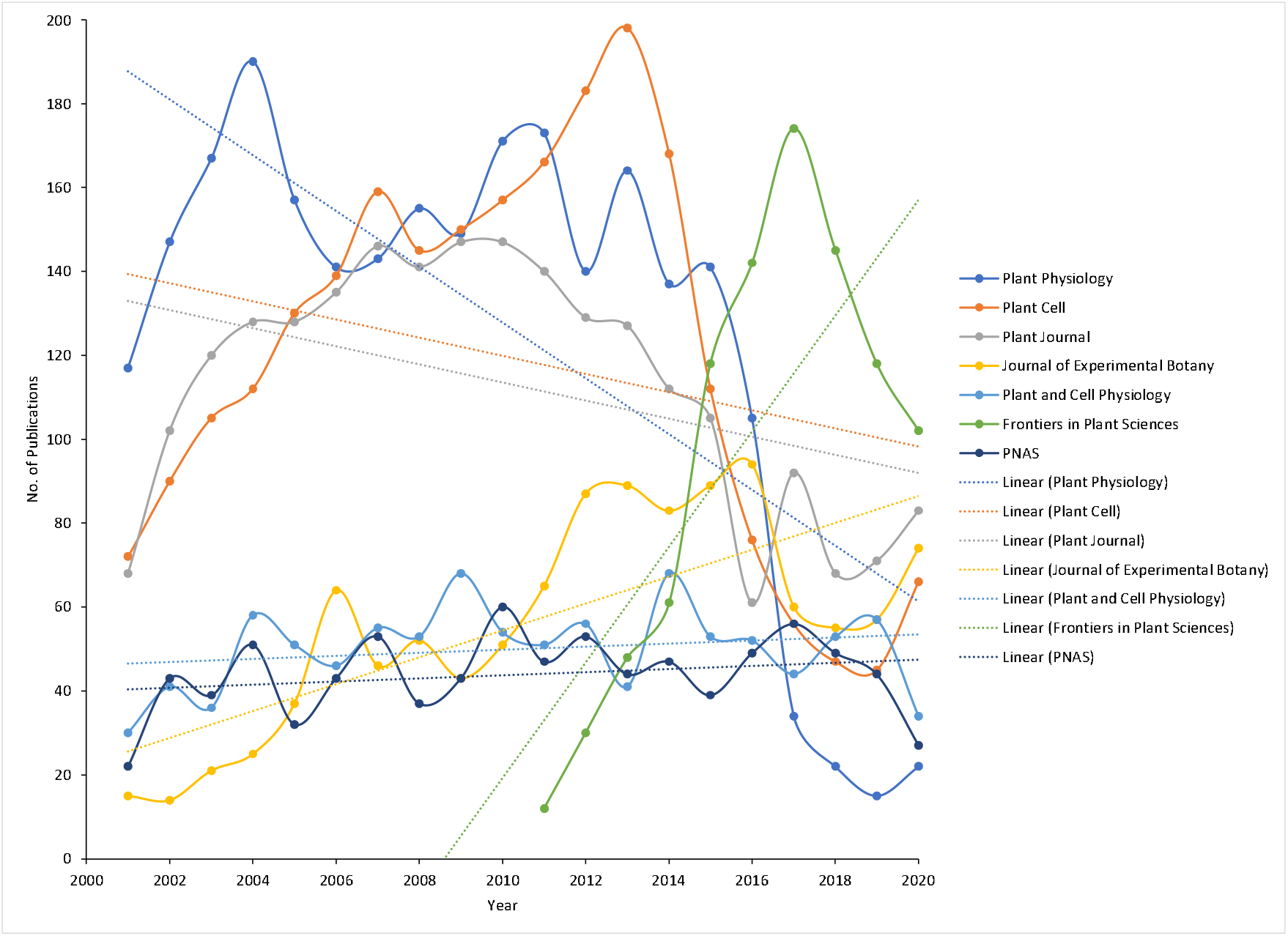
Performance of core journals (as defined by Bradford’s law) over 2001-2020 period based on total number of annual articles on ‘Arabidopsis’

## 5. Conclusion

We set out to determine the trajectory of Arabidopsis research in various areas. With the sequencing of the Arabidopsis genome, there was a rise in interest in the research, but it decreased after the advent and popularisation of new sequencing tools. Publication in Arabidopsis increased steadily in the first two quarters after the genome sequencing, reducing in the third quarter. Primary research interests spanned from mutant identification to protein investigation, as seen by co-word analysis, whereas gene expression experiments were constant throughout 20 years. Some of the highly cited articles were related to developing and validating novel techniques in plant science research.

Furthermore, Asian countries that have shown a strong interest in Arabidopsis research must collaborate closely with their western counterparts to improve the quality and reach. Quality is a significant consideration, as evidenced by the low number of citations, particularly from China, which has shown a rising trend in the total number of publications. The funds worldwide are depicted more toward the number of publications than ACPP. The target journal should be criteria based on impact factors for grants by the funding agency, depicting quality (IF) rather than quantity (T.P.). This can result in a good ACPP score, although ACCP is affected by several other factors also, viz., journal rank, primary/reporting author’s h-index, SCImago journal rank (SJR) and source normalised impact per paper (SNIP) (Yaminfirooz and Ardali, 2018). This can be represented from the author’s collaboration point of view as it also has a significant correlation with citation, and it has been witnessed that there is a lower collaboration among Asian countries compared to among European countries and USA. A substantial part of Arabidopsis papers has gone to three leading journals: Plant Physiology, Plant Cell, and Plant Journal. The growing trend toward open access publication is unmistakably the way forward for journals seeking to remain relevant.

